# Simultaneous Mapping of DNA Binding and Nucleosome Positioning with SpLiT-ChEC

**DOI:** 10.1101/2023.07.03.547581

**Authors:** Orion G. B. Banks, Michael J. Harms, Jeffrey. N. McKnight, Laura E. McKnight

## Abstract

The organization of chromatin – including the positions of nucleosomes and the binding of other proteins to DNA – helps define transcriptional profiles in eukaryotic organisms. While techniques like ChIP-Seq and MNase-Seq can map protein-DNA and nucleosome localization separately, assays designed to simultaneously capture nucleosome positions and protein-DNA interactions can produce a detailed picture of the chromatin landscape. Most assays that monitor chromatin organization and protein binding rely on antibodies, which often exhibit nonspecific binding, and/or the addition of bulky adducts to the DNA-binding protein being studied, which can affect their expression and activity. Here, we describe SpyCatcher Linked Targeting of Chromatin Endogenous Cleavage (SpLiT-ChEC), where a 13-amino acid SpyTag peptide, appended to a protein of interest, serves as a highly-specific targeting moiety for in situ enzymatic digestion. The SpyTag/SpyCatcher system forms a covalent bond, linking the target protein and a co-expressed MNase-SpyCatcher fusion construct. SpyTagged proteins are expressed from endogenous loci, whereas MNase-SpyCatcher expression is induced immediately before harvesting cultures. MNase is activated with high concentrations of calcium, which primarily digests DNA near target protein binding sites. By sequencing the DNA fragments released by targeted MNase digestion, we found that this method recovers information on protein binding and proximal nucleosome positioning. SpLiT-ChEC provides precise temporal control that we anticipate can be used to monitor chromatin under various conditions and at distinct points in the cell cycle.

## Introduction

The physiological state of a cell is largely defined by which genes are expressed at a given time. DNA-binding proteins (DBPs), particularly transcription factors (TFs) control which genes are expressed in all organisms.^1,2^ TFs bind to specific base sequences in the genome, thus targeting the transcriptional machinery to specific genes.^3,4^ Eukaryotes add another layer of regulation, controlling the accessibility of DBP binding sites via chromatin structure. Eukaryotic cells loop their DNA around histones, which then interact with one another to form chromatin.^5^ To a first approximation, the density of the chromatin structure controls accessibility: the denser the chromatin, the less accessible its TF binding sites and the less likely its genes will be transcribed.^6^

The interplay between DBP binding and chromatin structure is complicated and, despite decades of work, remains poorly understood. For example, in some circumstances, chromatin accessibility can be a poor predictor of transcription factor binding.^7,8^ DBPs can also promote chromatin remodeling, thus promoting the accessibility of specific genes.^9,10^ A complete understanding of transcriptional control in eukaryotes thus requires a deep understanding of the mechanisms that determine chromatin structure, the nature of DBP binding, and how these two processes interact.

One of the major challenges in picking apart the relative contributions of TFs and chromatin structure is a lack of tools that simultaneously probe the properties of chromatin and DBP binding. There are excellent tools to probe the locations of DBPs on the genome, including ChIP,^11^ ChIP-exo,^12^ and ORGANIC;^13^ however, these methods do not provide information about the chromatin environment surrounding the bound TFs. MNase digestion plays a complementary role, revealing chromatin structure by digesting any DNA not wrapped around histones.^14^ Several new techniques allow simultaneous assessment of chromatin state and DBP binding. These include CUT&RUN^15^ and CUT&Tag.^16^ While powerful, these techniques require complex setups, using antibody binding or transposases.

Antibody-free systems that capture DNA-protein interactions have previously been successful,^17^ including chromatin endogenous cleavage and sequencing (ChEC-Seq).^18,19^ ChEC-Seq employs a direct fusion of MNase to a protein of interest and maps cleavage sites adjacent to predicted consensus motifs. By relying on the inherent targeting capacity of TFs and transcription machinery, MNase digestion can be performed in situ without the need to select fragmentation sites via antibody recognition. However, directly fusing a large moiety like MNase, which is ∼17 kDa, to a protein can affect its expression and potentially its ability to bind DNA in the same manner as endogenous protein.^20^

Here, we present **<u>Sp</u>**yCatcher **<u>Li</u>**nked **<u>T</u>**argeting of **<u>Ch</u>**romatin **<u>E</u>**ndogenous **<u>C</u>**leavage combined with high-throughput sequencing (SpLiT-ChEC-Seq), a new technique which enables simultaneous mapping of protein binding signal and proximal nucleosome patterning at genomic sequences bound by endogenously expressed factors. This technique works without the requirement for antibody recognition or attachment of a large moiety to the DBP of interest. Instead, we use the SpyCatcher/SpyTag system^21,22^ to create inducible fusions of MNase with a target DBP. This allows us to precisely control when the MNase-DBP fusion is created, and separately, cleavage of nearby DNA by the nuclease. Further, we find that the resulting high-throughput output reports on both DBP binding and nucleosome positioning. Fragment size can be used to discern between nucleosome information and small DBP binding to reveal the target site and proximal nucleosome positions.^23,24^ We anticipate that this method will be useful for monitoring DBP binding and nearby nucleosomes simultaneously, with fine temporal control.

## Results

### Design of the SpLiT-ChEC System

We set out to create a system that would allow detection of protein-DNA interactions and nucleosome-derived signal, with no requirement for crosslinking or antibody recognition, by expressing a targetable MNase fusion in the yeast genome. MNase is suitable for this task because it requires millimolar concentrations of Ca^2+^ to be activated, which is several orders of magnitude higher than the typical Ca^2+^ concentration found in yeast cells.^25^

To target nuclease digestion, we utilized the SpyCatcher/SpyTag system.^21,22^ When co-expressed, SpyCatcher protein and SpyTag, a 13 amino acid sequence, form a covalent isopeptide bond, fusing together any protein attached to either component of the system.^21^ We fused MNase to Spycatcher via a flexible linker^26^ and placed it under an inducible promoter in the genome of S. cerevisiae (**Figure 1a**). Simultaneously, we introduce a SpyTag on a DBP at its endogenous locus (**Figure 1a**). This allows the SpyTagged-DBP to be expressed with addition of only a small polypeptide (**Figure 1b**). We anticipate that SpyTagged proteins exhibit native behavior, considering the diminished potential for misfolding or reduced binding capability imparted by a small, unstructured tag. The DBP therefore binds to its sites on DNA relatively unhindered, under the desired experimental conditions (i.e. cell cycle synchronization, transcription arrest, stress conditions, etc).

**Figure 1.**
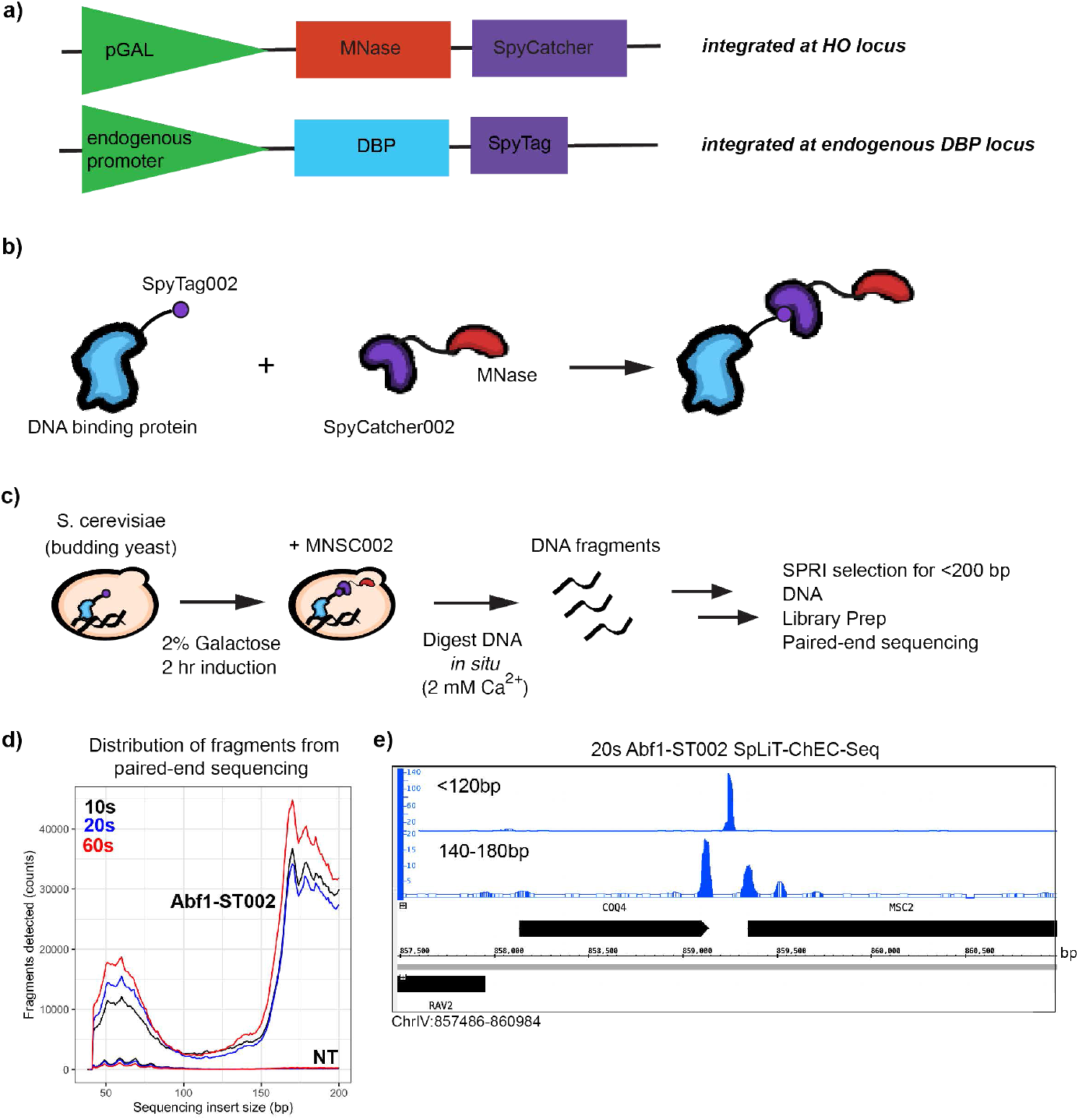
SpLiT-ChEC-Seq captures protein binding and nucleosome localization signal from sites of targeted nuclease digestion. **(a)** schematic of the constructs integrated into the genome for galactose-inducible MNase-SpyCatcher (*top*) and SpyTagged-factor of interest (*bottom*) **(b)** cartoon representation of the Spycatcher/Spytag targeting system employed to localize MNase to protein-bound loci in the yeast genome. **(c)** outline of the basic workflow to express MNase-Spycatcher and perform in situ digestion of DNA, which is purified and subjected to next-generation sequencing. **(d)** fragment size distributions observed across three timepoints of both untargeted (No-tag, NT) and Abf1-targeted SpLiT-ChEC-seq. Abf1 was C-terminally tagged with Spytag at the endogenous locus in yeast harboring the GAL-inducible MNase-Spycatcher construct. **(e)** Integrated Genome Browser (IGB) screen capture showing the signal over background for small (<120bp) and large (140-180bp) fragment protection observed in Abf1-targeted SpLiT-ChEC-seq after 20 sec. of calcium-activated DNA digestion.

Then expression of MNase-SpyCatcher (MNSC) is induced by galactose to create the DBP-SpyCatcher fusion (**Figure 1b**), and cleavage of DNA by MNase occurs only upon addition of Ca^2+^, giving temporal control to each aspect of the process (**Figure 1c**). By limiting expression of MNSC to a brief period prior to cell harvest, we observed low background digestion, presumably due to the difference in digestion rates that can be achieved by tethered vs untethered MNSC.

As a proof of concept, we tagged the protein ARS-binding factor 1 (Abf1) and performed SpLiT-ChEC at three time points after Ca^2+^ addition, which mirror the time points used to validate ChEC-Seq.^19^Abf1 presents an ideal model system to evaluate SpLiT-ChEC because of its known roles in chromatin organization, its activity as a GRF, and the well-defined consensus motif associated with its localization.^10,12,27,28^ MNSC expression was induced by addition of galactose, leading to formation of the Abf1-MNase fusion. After 2 hours of galactose induction, calcium was added to activate MNase, cleaving nearby DNA. We then extracted yeast genomic DNA, purified fragments <200 bp, and performed paired-end sequencing (**Figure 1c**).

The resulting technique, SpLiT-ChEC-Seq, enables simultaneous mapping of protein binding signal and proximal nucleosome patterning at genomic sequences bound by endogenously expressed factors.

### Abf1 SpLiT-ChEC produces well-defined signal with multiple fragment sources

The sequenced fragments from Abf1 SpLiT-ChEC showed a bimodal distribution of sizes: small protection factor fragments (SFP; < 120 bp) and large protection factor fragments (LFP; 140-180 bp) (**Figure 1d**). We hypothesized that fragmentation is influenced by protection of DNA associated with proteins (SFP) and nucleosomes (LFP). As a preliminary test of this hypothesis, we independently aligned the 120 bp and 140-180 bp fragments to the yeast genome. We found these fragments did, indeed, correspond to different sites on the genome (**Figure 1e**).

To quantify the results, we compared the 1x normalized genome coverage for fragments extracted from Abf1-tagged and non-tagged (“No-tag” or “NT”) samples at each timepoint to calculate the ratio of signal over background that occurs when MNSC is targeted to Abf1-bound loci. Both SFP and LFP fragments exhibited high-intensity peaks, primarily found in promoter and terminator regions of genes throughout the genome. When comparing the two signal sets, most often SFP signal is found in sequences lacking LFP signal, with LFP signal directly proximal to SFP sites at many targets, as is the case in **Figure 1e**.

In order to more directly compare our results to traditional methods such as ChIP-Seq and ChEC-Seq, we visualized the entire complement (<200bp) of fragments (**Supplemental Figure 1**). However, we found that separately aligning fragments from two size ranges allowed TF-derived small factor protection (SFP) (<120bp) to be viewed separately from the large fragment protection (LFP) (140-180bp), which possibly results from nucleosomes bound near Abf1-targets (**Figure 1e**).

### Canonical Abf1 binding and chromatin organization is captured in SpLiT-ChEC

To further test the hypothesis that SFPs probe transcription factor binding sites and LFPs probe chromatin organization, we asked how the SpLiT-ChEC signal mapped to the known Abf1 DNA binding motif (5’-WHWTCGTRTAWAGTGAYAND-3’, as determined by ChIP-exo).^12^ We examined our SpLiT-ChEC datasets and found many examples of Abf1 SFP directly situated over DNA sequences matching this motif (**Figure 2a**). To visualize SpLiT-ChEC signal at predicted Abf1 target regions in the yeast genome, we used FIMO^29^ to locate Abf1 consensus sequences based on the JASPAR30 entry corresponding to the Abf1 ChIP-exo footprint (MA0265.2, **Figure 2b**). We aligned and averaged our SpLiT-ChEC reads for all 2272 predicted Abf1 consensus sequences, setting the start of the consensus sequence to 0 bp.

**Figure 2.**
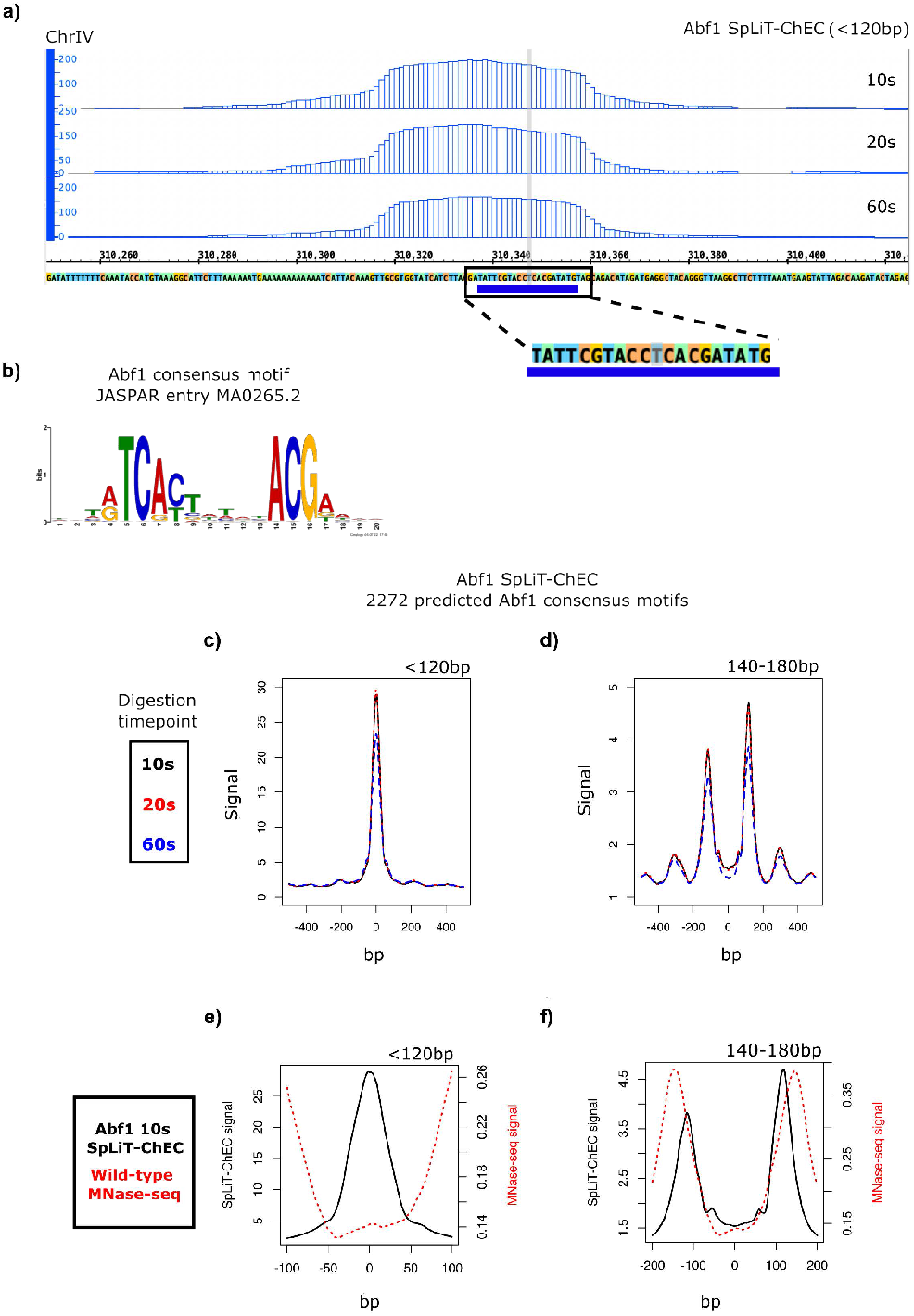
Abf1 SpLiT-ChEC signal is present at Abf1 consensus sequences and recapitulates chromatin organization elements observed in nucleosome positioning data. **(a)** IGB screen capture of Abf1 SpLiT-ChEC 10s, 20s, and 60s digestion timepoints showing small fragment signal located over an instance of the 20bp Abf1 consensus sequence (blue underline). **(b)** JASPAR entry MA0265.2, the 20 bp Abf1 consensus sequence derived from ChIP-exo. **(c)** Overlay of the average Abf1 SpLiT-ChEC small fragment signal from 10s, 20s, and 60s timepoints found at all 2272 Abf1 ChIP-exo consensus sequences in the yeast genome predicted by FIMO based on the motif found in (b). **(d)** Overlay of the average Abf1 SpLiT-ChEC large fragment signal from 10s, 20s, and 60s timepoints found at all 2272 predicted Abf1 ChIP-exo consensus sequences in the yeast genome. **(e)** Average traces of Abf1 10s SpLiT-ChEC small fragment signal and MNase-seq signal found at all 2272 predicted Abf1 ChIP-exo consensus sequences in the yeast genome. f) Average traces of Abf1 10s SpLiT-ChEC large fragment signal and MNase-seq signal found at all 2272 predicted Abf1 ChIP-exo consensus sequences in the yeast genome.

The average SFP signal was high directly over the Abf1 consensus motif, consistent with Abf1 binding at these sites and protecting them from digestion (**Figure 2c**). In contrast, the LFP signal was not typically found over consensus sequences (**Figure 2d**). Instead, it exhibits a series of flanking peaks whose intensity decays the further they are from the consensus sequence. The LFPs are thus revealing features besides TF binding.

We initially collected data from three sequenced timepoints (10s, 20s, 60s) in order to compare our data to that collected for Abf1 using ChEC-Seq^19^ for validation, and ensure that our system works on a similar timescale. We overlaid SpLiT-ChEC signal from all three datasets and showed that the shape and location of averaged SFP and LFP signal is similar across all timepoints (**Figure 2**). Time-dependent changes in signal intensity are present in both signal sets, most obviously seen as decreases in height of SFP and LFP dominant peaks in 60s timepoint traces (**Figure 2c, 2d**). This observation is consistent with data from ChEC-Seq, which can discern between high and low scoring pre-annotated binding sites based on time-dependent variability in cleavage intensity.^19^ While we do not see large differences across timepoints, we wished to demonstrate the ease of collecting temporal data, which may be useful for observing time-dependent processes such as nucleosome recovery and TF binding after transcription.

To test the hypothesis that the LFP signal reveals nucleosome positioning, we overlaid the SpLiT-ChEC SFP and LFP signals with nucleosome positions derived from MNase-Seq (GEO: GSE141676).^31^ MNase-Seq is used to map nucleosome positions in genomic DNA (**Supplemental Figure 2a**). As one would predict, the Abf1 SFP signal is present within the bounds of the nucleosome-free region (NFR) defined by MNase-Seq data - that is, it is not occluded by nucleosomes (**Figure 2e**). The LFP signal, by contrast, exhibited a striking similarity between the protected regions measured by MNase-Seq (**Figure 2f**).

The strong correlation between the SFP signal and the known consensus motif of ABf1 (**Figure 2c**), combined with the strong correlation between the LFP signal and nucleosome positions inferred by MNase-Seq (**Figure 2f**), leads us to conclude that SpLiT-ChEC provides information about both protein binding (SFP) and nucleosome positions (LFP) in a single measurement.

### Abf1 SFP can be used to find peaks with MACS3

We next sought to develop a method to use SpLiT-ChEC to identify binding targets throughout the genome. Analysis of aligned reads with MACS3 (https://macs3-project.github.io/MACS/) generated three sets of SFP-derived peaks of accumulated reads, one from each timepoint in Abf1-targeted SpLiT-ChEC. Variable numbers of peaks were called from each timepoint, with the highest number of peaks coming from the 20s signal (**Supplemental Figure 2b**). Rather than handling each timepoint peak set independently, we designed a computational process to identify target regions in Abf1 SFP signal according to peak groupings from across timepoints. We took the central two basepairs (dyads) from all peaks identified by MACS3 in each timepoint sample (n=4322); merging this set of dyads without extension resulted in 3476 independent entries. Next, the bounds of each target site were symmetrically extended by increasing values and any regions overlapping because of the expanded region size were merged. By visualizing the region counts, we anticipated that an optimal merge distance could be identified as the extension value where the total peak counts settled into a local minimum. A bar chart of the number of regions after each cycle showed substantial decreases at low merge distances, demonstrating that peaks were detected in similar regions across all timepoints (**Supplemental Figure 2c**). When we applied this methodology to Abf1-derived SFP peaks, our analysis suggested Abf1-targeted signal occurs within 90bp regions (n = 1807), which is consistent with the approximate size of the NFR present in MNase-seq and SpLiT-ChEC LFP (**Figure 2f**). Plotting SpLiT-ChEC signal using the calculated target sequences displays center-aligned SFP (**Supplemental Figure 2d**) and LFP (**Supplemental Figure 2e**), similar to that observed for signal at consensus sequences, suggesting our merge process identified the centers of Abf1-targeted regions from across all timepoints. We have performed SpLiT-ChEC on other DPBs and have observed that these time points are a good starting point to optimize the experiment, but we anticipate that different systems and/or DBPs may require collection of other timepoints depending on binding kinetics and the timing of the processes being studied.

### A minimal Abf1 motif marks canonical targets in SpLiT-ChEC-Seq

With Abf1 target regions in hand, we were curious if the Abf1 consensus motif was enriched in our recovered DNA sequences. Motif enrichment analysis using SEA (Simple enrichment analysis),^32^ which requires input of the Abf1 consensus motif to search data, showed that 46% (Q value = 5.69E-224) of the 1627 sequences from the input (randomly selected by SEA as a test set) contained a motif matching JASPAR entry MA0265.2. We also performed de novo motif discovery via XSTREME^33^ on 90bp sequences with Abf1 signal, which independently revealed a “minimal” version of the known Abf1 motif: 14bp vs the 20bp motif from ChIP-exo (JASPAR entry MA0265.2). This ‘minimal motif (MM) was found by SEA in 53% (Q value = 1.17E-250) of the sites (**Figure 3a**). Next, we calculated the number of Abf1 consensus motifs overlapped by Abf1 targets based on SpLiT-ChEC peaks, as determined by the extend-and-merge process. This analysis suggested Abf1-targeted SpLiT-ChEC digestion is found within 90bp regions. Plotting the fraction of FIMO-predicted motifs found in target regions from each merged peak set showed that the SpLiT-ChEC derived minimal motifs more frequently found in target centers (**Figure 3b**).

**Figure 3.**
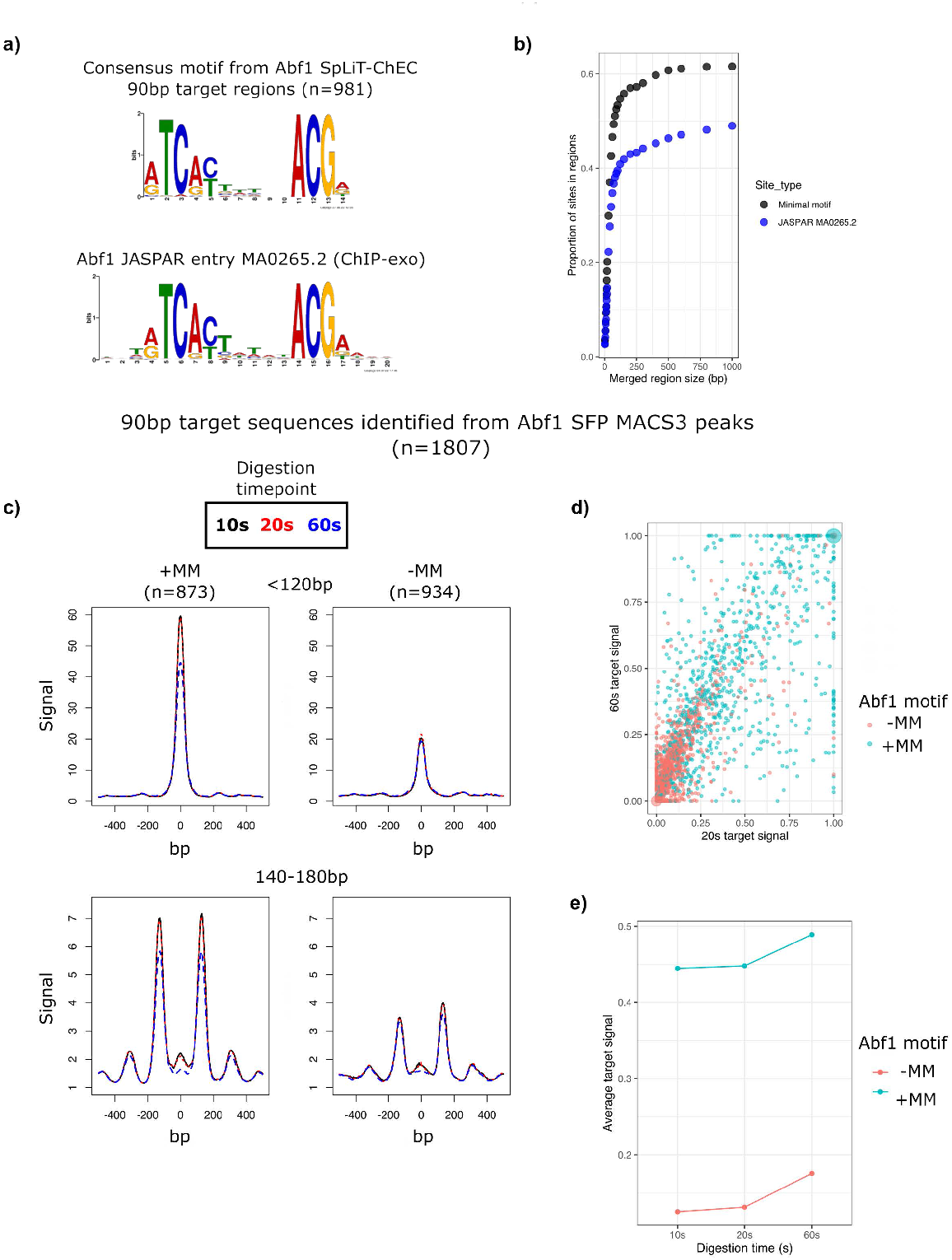
Abf1 SpLiT-ChEC allows consensus motifs to be identified and shows higher signal intensity within target regions containing consensus motifs. **(a)** Comparison of motifs identified for Abf1 from SpLiT-ChEC (top, 14bp) and ChIP-exo (20 bp, bottom). **(b)** Plot showing the proportions of 14bp SpLiT-ChEC minimal motifs (black) and 20bp ChIP-exo motifs (blue) found in regions identified by merging MACS3 peaks derived from Abf1 SpLiT-ChEC 10s, 20s, and 60s signal. **(c)** Overlay of average Abf1 SpLiT-ChEC signal centered on 90bp target sequences based on MACS3 peaks from 10s, 20s, and 60s timepoints. Signal is separated into sites containing a predicted 14bp minimal motif (+MM) and sites without a motif (-MM). **(d)** Per basepair Abf1 small fragment signal intensity found in 90bp target regions for 20s and 60s digestion timepoints, plotted as Cartesian coordinates and colored by sites with a minimal motif (+MM, blue) and without a minimal motif (-MM, orange). Scaling was achieved using the 5th and 95th percentile values for each timepoint as minimum and maximum values, respectively. Any values < 0 after scaling were assigned to 0 and values >1 were assigned to 1. **(e)** Abf1 small fragment signal intensity at all 90bp target sequences from all timepoints calculated as in (d), separated into +MM and -MM groups and averaged for each group and each timepoint.

Based on this result, we elected to use the FIMO-predicted MM sites as a filter to separate the 90bp target regions into +MM (with Minimal Motif) and -MM (without Minimal Motif) sets. Consistent with previous observations of lower perceivable levels of Abf1 binding at non-canonical sites,13 -MM target sequences showed overall lower average SFP and LFP (**Figure 3c**). Since the LFP shape does not appear to be disrupted at -MM sites, we presume that the lower LFP intensity observed at -MM sites is related to lower levels of Abf1-targeted digestion occurring in these regions, not less defined nucleosome patterning in the region. All of these-MM sites, which comprise 47% of total sites, contain motifs for the TF Reb1 and/or an ‘Abf1-like’ motif; this is comparable to the data from ORGANIC.^13^

Based on the observation that SpLiT-ChEC signal varies across time, we were curious if examining the per-basepair signal intensity within 90bp merge regions could be used to visualize the progress of MNase digestion as it occurs post-Ca^2+^ addition. Initially, we plotted two timepoints for each target region by treating the scaled intensity values as Cartesian coordinates (**Figure 3d**). Coloring the points based on the presence of Abf1 minimal motif (+/-MM) shows that there is separation between the two groups, with -MM generally showing lower intensity at both the 20s and 60s timepoints. Plotting average per-basepair intensities for each timepoint, separated by +/-MM, allowed us to view the trajectory of binding intensity over time (**Figure 3e**). Although the trends in SFP intensity appear similar, the overall intensity of signal without a minimal motif appears lower at all timepoints, consistent with observations made from plotting the raw SFP signal.

While the height of the peaks in SFP appear to decrease at 60s, the intensity per-basepair increases, which we anticipate is due to signal broadening as digestion progresses and DNA fragments are shortened.23 In summary, we observe a difference in the average Abf1 SpLiT-ChEC SFP signal intensity in target regions separated by the presence or absence of a minimal motif at each timepoint, where Abf1 targets without a minimal motif show lower average signal compared to sites that contain a minimal motif.

### SpLiT-ChEC signal can be clustered based on intensity and shape

Next, we wanted to see if unsupervised clustering of signal derived from SpLiT-ChEC would allow subsets of Abf1-targeted sequences to be identified. We applied k-means (k = 3) clustering to raw LFP signal from the 60s timepoint at 90bp target regions (**Supplemental Figure 3a**). The patterns observed in each LFP cluster were surprisingly distinct, with obvious signal bias towards either side of the Abf1 target center (clusters 1 and 2), and the 3rd cluster showing targets with LFP signal in the center of sites. Focusing on the central 200bp of the 90bp target regions (**Supplemental Figure 3b**) shows the asymmetry in LFP protection, with the most substantial LFP at the edges of the central 90bp in each target. When we plotted the SFP average signal trajectories for each LFP cluster (**Supplemental Figure 3c**), we observed similar signal evolution patterns for clusters 1 and 2, which remained mostly constant across timepoints.

Interestingly, the cluster 3 average trajectory shows lower intensities at each timepoint when compared to cluster 1 and 2 patterns, with a consistent upward trend in signal evolution. When k-means (k = 3) clustering was applied to SFP per-basepair intensity measured across the three timepoints, plotting the values for each site at 20s and 60s post-Ca^2+^ addition shows distinct groups of points (**Supplemental Figure 3d**). The values of average intensity per bases from each cluster suggest that k-means clustering separates sites primarily by signal intensity at each timepoint (**Supplemental Figure 3e**).

Plotting average LFP signal according to SFP cluster annotations (**Supplemental Figure 3f**) revealed signal shapes that are distinct from those found in LFP-based clusters (**Supplemental Figure 3a**). Unlike the LFP clusters, signal intensity on either side of the center within each cluster was roughly equal with no clear preference for sites with or without LFP signal in the central NFR between groups. Instead, LFP patterns associated with SFP clusters differ primarily by intensity, suggesting that the per-base SFP calculations are limited to the relative amount of Abf1-directed digestion occurring within the 90bp targets.

### Abf1 SpLiT-ChEC signal near TSSs is directional and highly variable

Given the directionality we observed in Abf1 LFP signal clusters (**Supplemental Figure 3**), we applied strand information derived from TSSs near 90bp targets to see whether signal is orientated with respect to the direction of transcription. Using T-Gene,^34^ we located TSSs in the budding yeast genome nearest to each 90bp target, limited to within +/-1000bp (n = 1660). After applying the strand designation for the identified TSSs, we plotted the average SFP signal at target centers across all timepoints (**Figure 4a**). The average traces of Abf1 SFP near TSSs show the expected high intensity central peak, but also a more prominent low-intensity protection peak roughly 200-250 bp upstream of Abf1 target centers, which was present on either side of the central peak before the addition of strand information. When we examined LFP signal at the TSS-proximal targets (**Figure 4b**), we noticed a prominent bias in peak intensity opposite from the direction of transcription at all timepoints. The relative peak intensity is in opposite orientation compared to +1/-1 nucleosome positions observed by MNase-Seq at NFRs near TSSs. While we cannot rule this out as a physical limitation of the SpLiT-ChEC system, this could also represent the presence of co-bound factors at Abf1 targets that help define precise nucleosome positions.^35,36^

**Figure 4.**
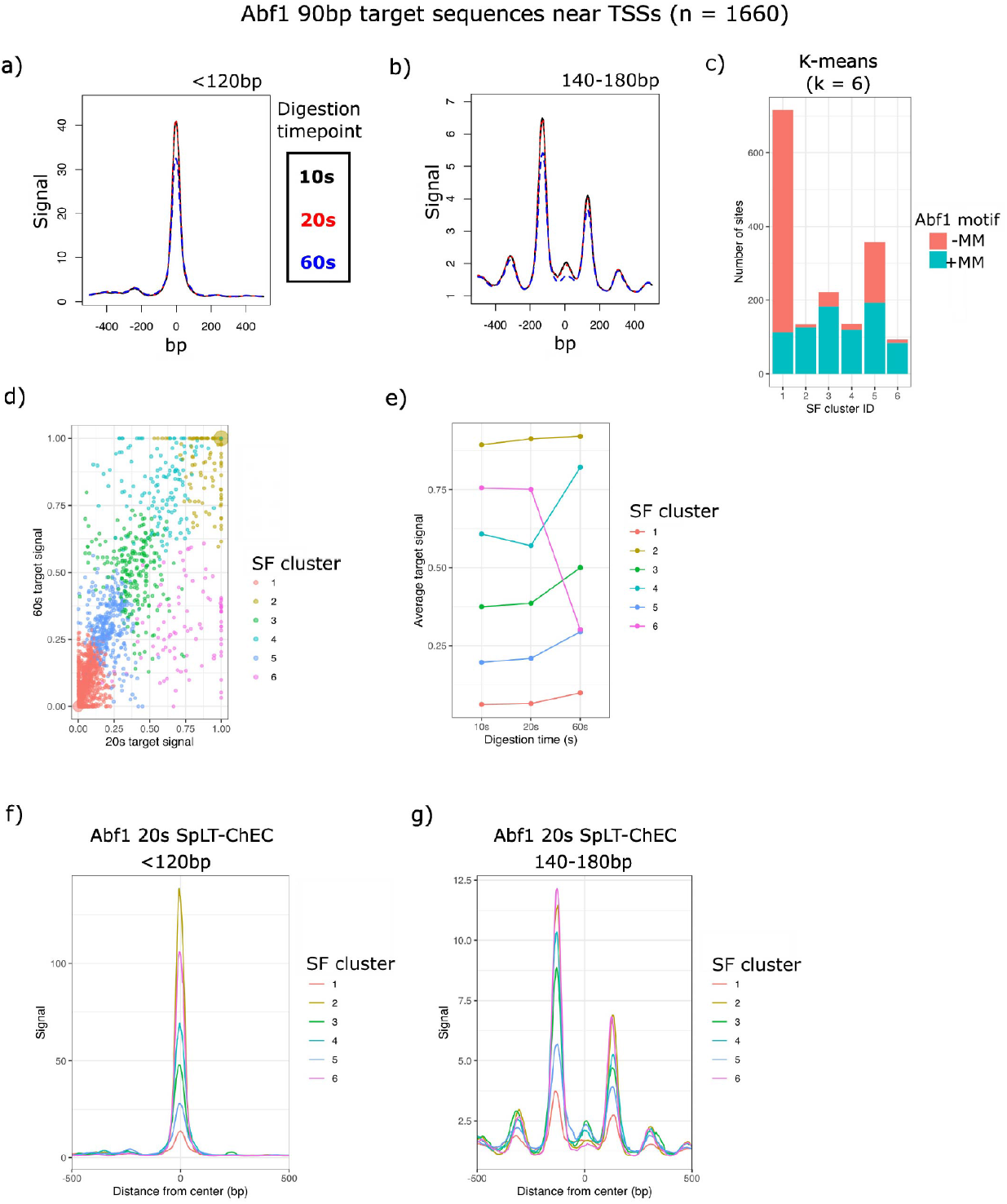
Abf1 SpLiT-ChEC small fragment signal near transcription start sites is directional with respect to gene orientation and can be used to identify similarly protected target sequences. **(a)** Overlay plot of 10s, 20s, and 60s Abf1 SpLiT-ChEC small fragment signal centered on 90bp targets within +/-1000 bp of a TSS. TSS strand designation was applied to plotting calculations. **(b)** equivalent to (a), except using long fragment signal. **(c)** Counts of the number of Abf1 90bp target sequences in each k-means cluster (k = 6) colored by sites with and without a minimal motif. **(d)** Per basepair Abf1 small fragment signal intensity found in 90bp target regions for 20s and 60s digestion timepoints, plotted as Cartesian coordinates and colored by k-means SF cluster designation. **(e)** Abf1 small fragment signal intensity at TSS-proximal 90bp target sequences from all timepoints calculated as in (d), separated into SF cluster groups and averaged for each group at each timepoint. **(f)** Average traces of 20s Abf1 SpLiT-ChEC small fragment signal separated by k-means cluster annotations. **(g)** equivalent to (f), except applied to 20s Abf1 long fragment signal.

We were curious if Abf1 SFP per-base intensity across timepoints for TSS-proximal targets would reveal subgroups based on optimized k-means clustering. We applied the WSS (within sum of squares) calculation to create an elbow plot (**Supplemental Figure 4**) and identified k=6 as an appropriate number of k-means clusters. To our surprise, each cluster contained both -MM and +MM sites (**Figure 4c**). While cluster 1 is seemingly dominated by -MM targets, no group appears to be completely composed of -MM or +MM sites.

Plotting the per-basepair SFP signal at 20s and 60s for each target site shows visually distinct groups of points throughout the coordinate plane when colored by cluster (**Figure 4d**). Using the SF cluster annotations, we plotted the average per base intensities at each timepoint (**Figure 4e**), revealing that the clusters are mostly separated by signal intensity, which was expected based on our preliminary cluster analysis (**Supplemental Figure 3**). However, some degree of the change in signal between timepoints appears to be captured within the k=6 clusters, particularly when examining clusters 4 and 6, which are graphically similar in intensity and trend for 10s and 20s, but display opposite trends when examining the 60s average per base signal. To examine the average signal identified within the groups of Abf1 targets, we plotted SFP (**Figure 4f**) and LFP (**Figure 4g**) according to SFP cluster labels, using only the average signal derived from the 20s Abf1 SpLiT-ChEC timepoint. The SFP signal traces differ based on central peak intensity, with some small differences in proximal signal, though no clear differences are readily identifiable. No obvious LFP patterning trends are evident in the average signal within the SF clusters, though each trace differs by overall intensity, matching expectations set by our pre-TSS cluster analysis. We anticipate this analysis could be more insightful for proteins directly involved in nucleosome positioning, which may show distinct LFP signal patterns associated with targeted chromatin remodeling.

### Abf1 SpLiT-ChEC-seq LFP near TSSs reveals distinct clusters of chromatin organization

We wanted to visualize the differences in LFP patterning among 90bp targets near TSSs, considering the known relationship between Abf1 localization, nucleosome positioning, and NFR formation at Abf1-targeted genome loci.10,27 We selected k = 8 as the optimal number of clusters, based on an elbow plot using the WSS method (**Supplemental Figure 5**). Applying k-means clustering to the signal from a single LFP timepoint, 60s, shows clusters that contain various proportions of sites both with and without an Abf1 minimal motif (**Figure 5a**); as with the SFP signal (**Figure 4c**), +MM and -MM sites do not separate into distinct clusters based on the presence or absence of a minimal motif. When we plotted the average 60s SpLiT-ChEC LFP signal based on long fragment (LF) cluster annotations, we observed differences in signal intensity, directionality, and the locations of the most intense LFP signal (**Figure 5b,c**). Similar to our preliminary clustering analysis, we observe a subset of sites with LFP signal in the predicted NFR. LF clusters 4 and 7 both display this feature, with cluster 7 showing a clear LFP peak in the center of the average signal traces (**Figure 5c**). The average LFP signal associated with cluster 6 shows a narrowing of the NFR with respect to other signal traces, an interesting feature which is often associated with decreased transcriptional activity for nearby target genes.^37^

**Figure 5.**
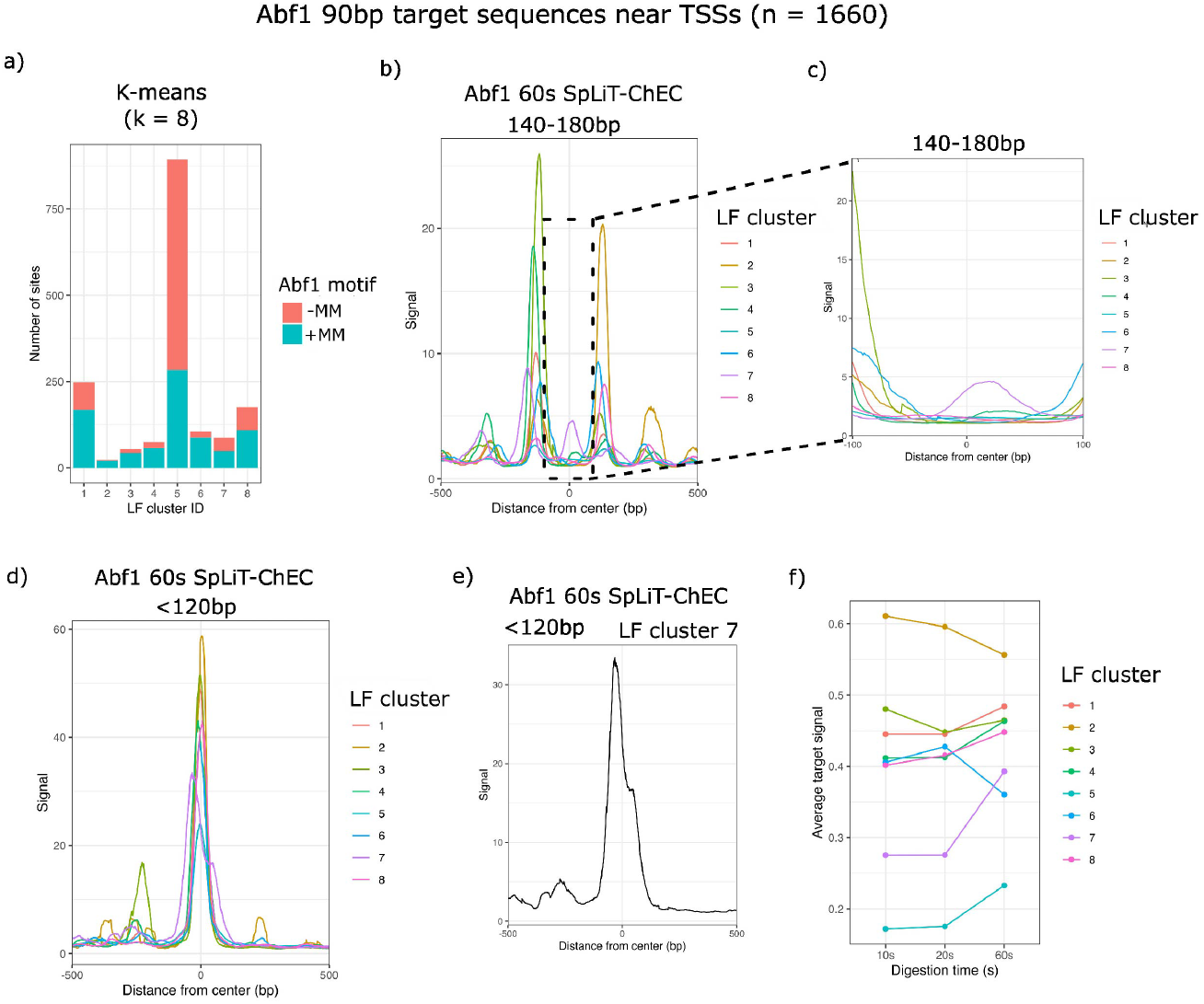
Abf1 SpLiT-ChEC large fragment signal near transcription start sites can be used to identify chromatin organization subgroups with distinct protection characteristics. **(a)** Counts of the number of Abf1 90bp target sequences in each k-means cluster (k = 8) colored by sites with and without a minimal motif. **(b)** Average traces of 60s Abf1 SpLiT-ChEC large fragment signal separated by k-means cluster annotations. **(c)** Close up view of the signal within the central 200bp of traces found in (b). **(d)** equivalent to (b), except plotting 60s Abf1 SpLiT-ChEC small fragment signal. **(e)** trace showing the average small fragment signal of target sites found in LF cluster 7, which contains a centralized region of long fragment protection. **(f)** Abf1 small fragment signal intensity at TSS-proximal 90bp target sequences from all timepoints, separated into LF cluster groups and averaged for each group at each timepoint.

We were curious about the relationship, if any, between the differences we observed in average LFP signal among LF clusters and the corresponding SFP signal qualities. We were surprised to find that SFP signal patterns in these clusters vary substantially in signal intensity and shape, with various degrees of SFP outside of the expected 90bp central region (**Figure 5d**). While every average SFP signal trace appears to show some amount of protection proximal to the central high-intensity signal, cluster 3 provides the most striking example. Cluster 3 displays a defined peak 200-250bp upstream of the central SFP and contains a high intensity central SFP peak. The upstream signal feature originally became more pronounced when strand information was applied to TSS-proximal 90bp target sequences (**Figure 4a**). Interestingly, when examining the average LFP signal for cluster 3, we observed a high level of signal directly between the region defined by the two SFP average signal features. Additionally, we observed cluster 7 as having the most visually distinct average SFP signal (**Figure 5e**), which shows a broad shoulder in place of the high intensity, tightly defined signal we observe in other clusters. When compared to the average LFP trace for cluster 7 (**Figure 5c**), this suggests that centralized LFP signal is not an effective predictor of Abf1 localization, but rather that Abf1 is capable of targeting digestion to these sites regardless of LFP within the expected target regions. Further, when we examined the average per-base SFP intensity at each timepoint for the LF-derived clusters (**Figure 5f**), we found several interesting patterns, most notably in clusters 5 and 2. Cluster 5 displays the lowest overall intensities among this set of clusters, which we anticipate could be related to the relatively high number of target sequences in this group that lack an Abf1 consensus motif (**Figure 5a**). Cluster 2 essentially displays the opposite effect – the relative number of -MM sites is lower for this group of Abf1 targets when compared to cluster 5 and average per base SFP intensity appears to be higher at all timepoints.

## Discussion

While many methods exist that allow protein-DNA interactions to be detected, few allow transcription factor binding sites and chromatin organizational elements to be measured simultaneously. Generally, multiple experiments must be performed if information on both protein-DNA interactions and nucleosome positions is required to make conclusions about the behavior of proteins related to chromatin organization. Further, current methods for producing genomic DNA fragments from multiple sources are either non-targeted (ATAC-Seq, MNase-Seq) or require preparation of functionalized antibodies to target enzymatic digestion (CUT&RUN) or transposase activity (CUT&Tag) to loci of interest. SpLiT-ChEC provides an alternative to these systems, as it relies on the inherent capacity for SpyCatcher and SpyTag to self-associate in a biological environment. By SpyTagging proteins in the yeast genome, expression is controlled by an endogenous promoter, which avoids the issue of mismatched expression associated with the use of a plasmid and/or exogenous promoter. Since the SpyTag is small (13 amino acids), it is straightforward and inexpensive to add to an endogenous protein in a variety of organisms and cell types.

Integrating MNSC into the yeast genome with an inducible promoter provides two benefits: first, a common base strain can be used as a platform for SpyTagging proteins of interest and second, inducing MNSC expression for only a short period before harvesting cultures allows the SpyTagged target protein to be expressed in its near-native state and bind to chromatin without interference from a large fusion protein. After expressing the MNSC construct, MNase is covalently tethered to the SpyTagged target and DNA digestion is activated by adding high concentrations of Ca^2+^ to permeabilized yeast cells. This provides better control of MNase activity, which reduces the level digestion by free MNSC and provides high signal-over-background in our calculated coverage. By separating the sequenced DNA fragments into predefined size ranges, we assess the levels of protection from DNA digestion corresponding to DNA binding proteins (<120bp fragments) and nucleosomes (140-180bp) from a single paired-end sequencing run. Our results suggest that SpLiT-ChEC can serve as an effective tool for monitoring protein localization and proximal nucleosome patterning at DNA sequences that are bound by SpyTagged proteins.

To test our system, we chose Abf1, an important regulator of chromatin structure that helps establish NFRs at loci throughout the yeast genome.10,27 NFRs are associated with GRF binding and are frequently found surrounding consensus motifs,28 which was readily observed in the LFP signal we detected with SpLiT-ChEC. When compared to SFP signal, LFP signal is generally found outside of regions associated with protein binding, like those marked by Abf1 consensus sequences. This aligns with the observation that Abf1 can nucleate NFRs by competing with nucleosomes to bind DNA, similar to the activity observed for pioneering factors in higher eukaryotes.27,38 While Abf1 is not capable of repositioning nucleosomes, GRF-mediated targeting of chromatin remodeling proteins, like RSC, is a known mechanism that allows nucleosomes to be evicted from NFRs throughout the genome.35,36,39 Consistent with this activity is the appearance of well-defined LFP signal in many of the regions directly proximal to the Abf1 target sites, which we defined using peak centers from multiple timepoints of SpLiT-ChEC SFP signal.

We predict that analyzing clusters of targets defined by SpLiT-ChEC could identify enriched sequence repeats, subsets of sequence motifs, or protein co-localization, any of which may correspond to biologically relevant mechanisms that define precise, site-specific nucleosome positions or enable association of factors with nucleosome-bound DNA. Based on our results, we anticipate SpLiT-ChEC will be a valuable tool for studying proteins that are directly involved in dynamic processes such as chromatin remodeling and transcription.

We have also collected SpLiT-ChEC data for Reb1, Med14, Ume6, and Gal4 (unpublished), demonstrating that this technique can be applied to a wide variety of chromatin-associated factors. Full analysis of this data was beyond the scope of this manuscript, but preliminary results suggest that SpLiT-ChEC is a flexible tool for detecting DNA binding and nucleosome patterning in each of these contexts.

When separating the Abf1 SpLiT-ChEC signal by the presence of a minimal consensus motif, we were surprised to see the relative amount of LFP definition was not obviously perturbed in sites lacking a predicted MM. While we cannot rule out that Abf1 is able to stimulate NFR formation at these sites, though with seemingly lower binding intensity, we imagine these interactions represent either co-targeting or binding to NFRs established by other GRFs found in yeast.^13,28,40^ We noted that, of the 1807 90bp sequences identified as Abf1 targets in this study, 296 also contain a consensus motif (JASPAR entry MA0363.2) associated with binding of the well-studied GRF Reb1,41 based on sites located in the yeast genome with FIMO pattern matching. However, after separating Abf1 targets into -MM and +MM groups, we found that 241 of those 296 Abf1 target sites do not contain an Abf1 consensus sequence. While a more thorough analysis is warranted to substantiate our hypothesis, we believe this highlights a valuable insight from SpLiT-ChEC data: Abf1 binding throughout the genome can occur independently of its consensus motif, but is not necessarily non-specific, as it appears to be associated with gaps in LFP signal, and, theoretically, specific chromatin organizational states that allow access to a subset of loci.

Since SpLiT-ChEC sequencing data is relatively complex to process and interpret, we invested substantial effort into designing computational pipelines and tools that simplify the analysis of our datasets, which is freely available on GitHub (Bankso/SCAR and Bankso/SEAPE). For example, the process we designed to merge peaks into high-confidence regions of Abf1 binding could be applied to SpLiT-ChEC data recovered from other factors, regardless of their association with NFRs. In the future, we predict that timepoints collected earlier after that addition of Ca2+ may be better suited to this task, as we observed a time-dependent shift in the population of large fragments, associated with extended MNase digestion, which could overrepresent SFP signal found in the experiments. As more SpLiT-ChEC data is generated, we hope to refine the computational processes supporting this analysis, with the intention to continue to generalize the approach and make it and easily accessible for studying chromatin organization.

## Materials and Methods

### Strain construction

Stock strains stored at -80°C were streaked onto selective media. From overnight starter cultures, yeast were diluted and grown at 30°C in yeast peptone media containing 2% glucose (YPD), then harvested by centrifugation at 3428 RCF for 5 minutes. Pellets were resuspended in sterile water and aliquoted to sterile tubes, then washed twice with 1 mL of sterile water. Washed pellets were resuspended in transformation buffer (240 uL PEG 50%, 36 uL LiOAc, 50 uL salmon sperm DNA) with a DNA cassette to be integrated into the genome via homologous recombination. Samples were placed at 42°C for a minimum of 1 hr before plating on YPD. Replica plates of transformants were made onto selective media after sufficient growth.

### Plasmid cloning

Gibson cloning^42^ was used to subclone SpyCatcher002 from Addgene 102827. Sticky-end PCR^43^ was performed as previously described to insert MNase-GGSx5-SpyCatcher002 into an HO-pGAL plasmid derived from Addgene 51664. The SpyTag002 plasmid was made by site-directed mutagenesis of a SpyTag001 plasmid, which was constructed via Gibson cloning using gBlocks from IDT.

### Yeast strain verification

Selection plates were grown at 30°C and screened for colonies containing integrated DNA at the required locus via colony PCR. With a toothpick, a small sample was taken from each candidate colony on the selective plate and stirred into 100 uL of lysis buffer, then heated at 65°C for 5 minutes. Next, 300 uL of EtOH were added and the samples were centrifuged for 5 minutes at 21130 RCF. After removing the supernatant, the pellets were washed with 300 uL 70% EtOH and allowed to dry at room temperature. Dried pellets were resuspended in 100 uL sterile water and centrifuged for 5 minutes at 21130 RCF, after which 50 uL of supernatant containing gDNA was transferred to new tubes. Genomic DNA samples were analyzed by PCR with sample-specific primers designed to amplify target regions of the genome where the integration was expected to occur. PCR reactions were analyzed by agarose gel electrophoresis. Positive transformants were streaked onto selection media and verified on YPG before use in experiments.

### Growth, harvest, and preparation of yeast for SpLiT-ChEC

Adapted from Zentner et al., 2015.^19^From overnight starter cultures, yeast were diluted, then grown in yeast peptone media containing 2% raffinose (YPR) at 30°C before adding 20% galactose to 2% final concentration. Cultures were returned to 30°C for 2 hours to express MNase-SpyCatcher. Next, cultures were centrifuged at 1500 RCF for 1 minute at room temperature. Pellets were resuspended in 1 mL buffer A (15mM Tris pH 7.5, 80 mM KCl, 0.1 mM EGTA, 0.5 mM spermidine, 1x Proteoloc protease inhibitors, 1x PMSF/ leupeptin/benzamidine) and transferred to 1.5 mL tubes. Samples were pelleted at 1500 RCF for 30 seconds and the supernatant was removed. This was repeated twice more with 1 mL buffer A. Washed pellets were resuspended in 570 uL of buffer A, then combined with 30 uL of 2% digitonin. Samples were briefly mixed and allowed to incubate at 30°C for 5 minutes to permeabilize cells. Permeabilized yeast were mixed again, then a 100 uL pre-digestion sample was removed to one of 6 tubes containing 10 uL 10% SDS and 90 uL of 2x ChEC stop buffer (400 mM NaCl, 20 mM EDTA, 4 mM EGTA).

### SpLiT-ChEC targeted DNA digestion

Adapted from Zentner et al., 2015.^19^To perform DNA digestion, 1 uL of 1 M CaCl_2_ (2 mM final) was added to the remaining permeabilized yeast, followed by briefly vortexing and placing the sample at 30°C, at which point a timer was started. Samples were collected at five time points post-Ca^2+^ addition by removing 100 uL of sample to pre-filled and labeled tubes with ChEC stop buffer and 10% SDS. All tubes were mixed immediately after adding a sample aliquot to ensure MNase digestion was deactivated. After collecting the final timepoint sample, 4 uL of 20 mg/mL proteinase K was added to each tube, followed by incubation at 55°C for a minimum of 30 minutes. At this point, samples were stored at -20°C or directly purified.

### DNA purification and size selection

Adapted from Zentner et al., 2015.^19^SpLiT-ChEC DNA fragments were isolated from proteinase K-treated samples by first adding 200 uL of phenol-chloroform-isoamyl alcohol (25:24:1) and vortexing thoroughly. Samples were centrifuged at 21130 RCF for 5 minutes, after which ∼180 uL of the aqueous (top) layer was removed to new 1.5 mL tubes. To precipitate DNA, 500 uL EtOH, 15 uL 3M NaOAc pH 5.3, and 1.5 uL (30 ug) of 10 mg/mL glycogen were added to each sample and placed at -80°C for at least 20 minutes. DNA pellets were resuspended in RNase solution (1 uL RNase A 20 mg/mL, 29 uL 1x Cutsmart or low-TE buffer) and digested at 37°C for at least 30 minutes. 5 uL of each timepoint sample was analyzed by agarose gel electrophoresis to confirm time-dependent DNA streaking. Three samples from across the timepoint range were used in size-selection with a 3:1 ratio of SPRI beads (AmpureXP). Supernatants from size selection were retained for each sample, primarily containing fragments <200 bp in size. The size-selected samples were PCI extracted and precipitated as above. The final DNA pellet was solubilized in 12 uL low-TE buffer and the concentration determined by Pico Green analysis (Thermo).

### SpLiT-ChEC NGS library preparation

For library preparation (NuGen Ultralow V2 kit), at most 10 ng of DNA was used from each sample as input and all steps were performed per the manufacturer’s protocol, except for the noted exceptions here. Library SPRI purification steps were performed with1.8x vol of beads to increase the small fragment content of the final libraries and 15 uL of low-TE buffer were used to elute final libraries, collecting 12 uL as a final sample for sequencing. Amplification cycles were scaled as necessary depending on DNA available, with 13 cycles used for 10ng inputs. Libraries were sequenced in paired-end mode for 37 cycles on an Illumina HiSeq 5000 system in high output mode.

### Data analysis

Paired-end reads received in FASTQ format were run through FastQC,44 then aligned to the Ensembl R64^45^ reference genome using bowtie2^46^ with the “--no-unal”, “--no-mixed” and “--no-discordant” flags. Fragment filtering to analyze small (0-120 bp), large (140-180 bp) and full (0-200 bp) fragment sets was accomplished with bowtie2 flags ‘-I’ and ‘-X’. Aligned reads were filtered (MAPQ > 30) and indexed with samtools.^47^ Deeptools^48^ ‘bamCoverage’ was used to determine sequencing coverage from filtered BAM files and output bigWig-formatted files were normalized to 1x coverage (RPGC, genome size of 12.1 megabases). Coverage files were processed with deeptools ‘bigWigCompare’ to calculate the ratio of sample coverage over background. Peaks were identified from sample BAM files by MACS3 (https://macs3-project.github.io/MACS/) ‘callpeak’ using the corresponding timepoint control BAM (“No tag”) as background with the flags ‘--keep-dup -all’ and ‘-f BAMPE’. Bedtools^49^ ‘intersect’ and ‘merge’ were used to subset/combine BED file entries. A pipeline was created to locate groups of peak centers from across timepoints using custom tools in python and functions from bedtools. Deeptools ‘computeMatrix’, ‘plotHeatmap’, and ‘plotProfile’ were used to generate matrices, heatmaps and average profiles according to input BED regions, respectively.

Predicted binding sites in the Ensembl reference genome for S. cerevisiae were located using FIMO^29^ with the standard cutoff score of 10-4. Published motifs were acquired from JASPAR 2022 database.^30^

To perform motif analysis, BED files containing target regions were converted to FASTA using bedtools ‘getfasta’. Output FASTA files containing target sequences, now fragments of the yeast genome, were processed with MEME suite XSTREME^33^ using standard input parameters and the JASPAR 2022 non-redundant database, except the motif range was expanded to 5-20bp. This also produces a list of motifs found in the submitted FASTA fragments, which can be viewed in **Supplementary Tables 1 and 2**. The most enriched motifs that grouped with JASPAR entries for tagged DBPs were used with FIMO (standard cutoff) to locate sites in the yeast genome.

All scripts and tools used to analyze SpLiT-ChEC data are available on GitHub (Bankso/SCAR and Bankso/SEAPE).

## Author Contributions

Conceptualization, O.G.B.B., L.E.M., J.N.M.; Methodology, O.G.B.B., M.J.H, L.E.M., J.N.M.; Investigation, O.G.B.B, L.E.M., J.N.M.; Additional Resources, M.J.H.; Writing - Original Draft, O.G.B.B.; Writing - Reviewing and Editing, O.G.B.B., M.J.H., L.E.M.; Visualization, O.G.B.B., M.J.H, L.E.M.; Supervision, M.J.H., J.N.M., L.E.M.; Project Administration, L.E.M., J.N.M.; Funding Acquisition, J.N.M.

## Data Availability

The datasets generated during this study are available at GEO under accession code GSE236273.

## Acknowledgments

The authors thank the Genomics Core (GC3F) at the University of Oregon for high throughput sequencing services. We would also like to thank Gabe Zentner for helpful discussions regarding experimental design and data analysis. This work was supported by a National Institutes of Health training grant T32 GM007759 (to O.G.B.B), and by NIGMS R01 GM129242 (to J.N.M. and L.E.M).

## Supplemental Figures

**Supplemental Figure 1.**
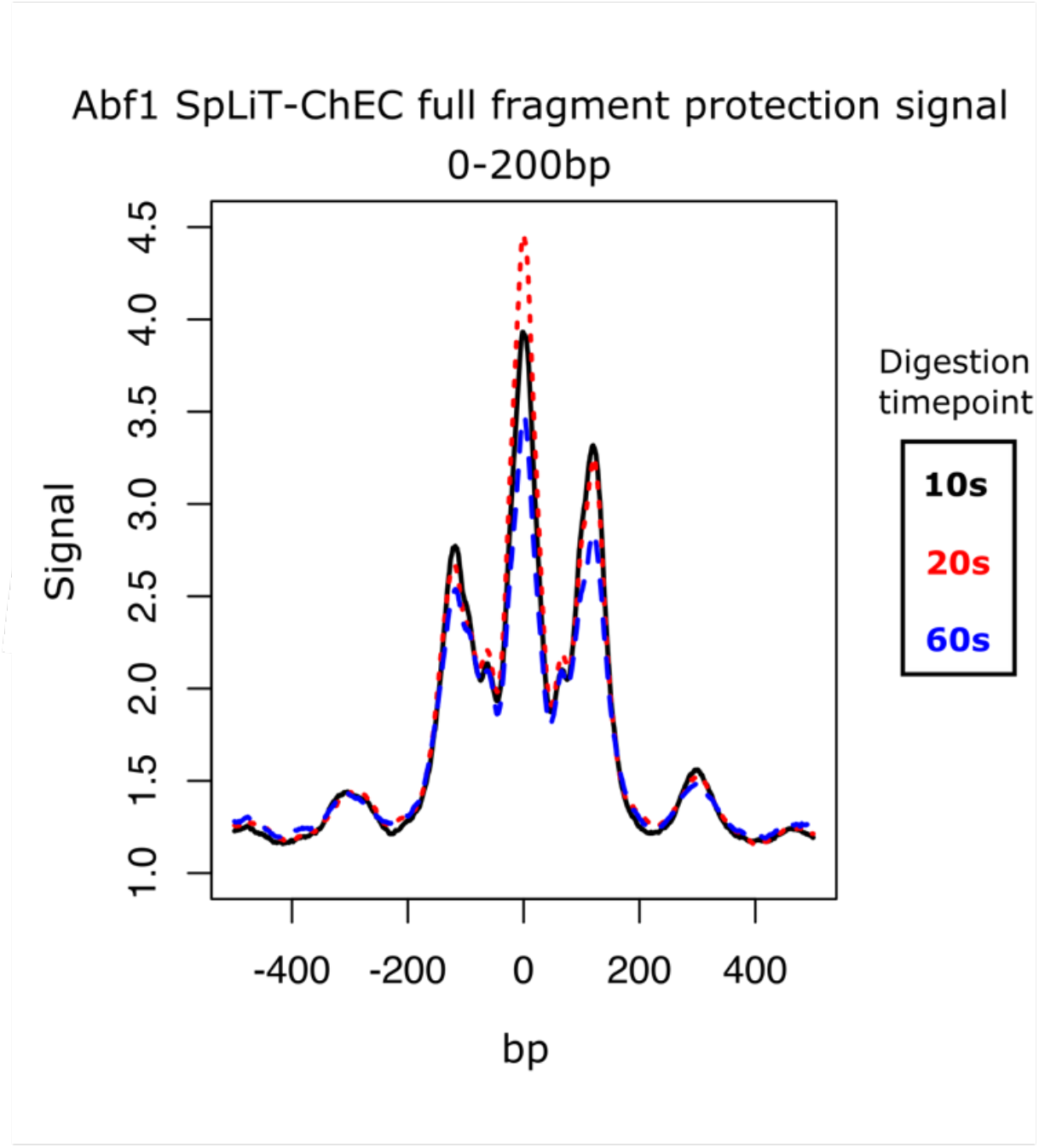
Abf1 SpLiT-ChEC total fragment pattern is a composite of SFP and LFP signal. Overlay of signal derived from all DNA fragments less than 200bp collected at three timepoints of Abf1-targeted SpLiT-ChEC

**Supplemental Figure 2.**
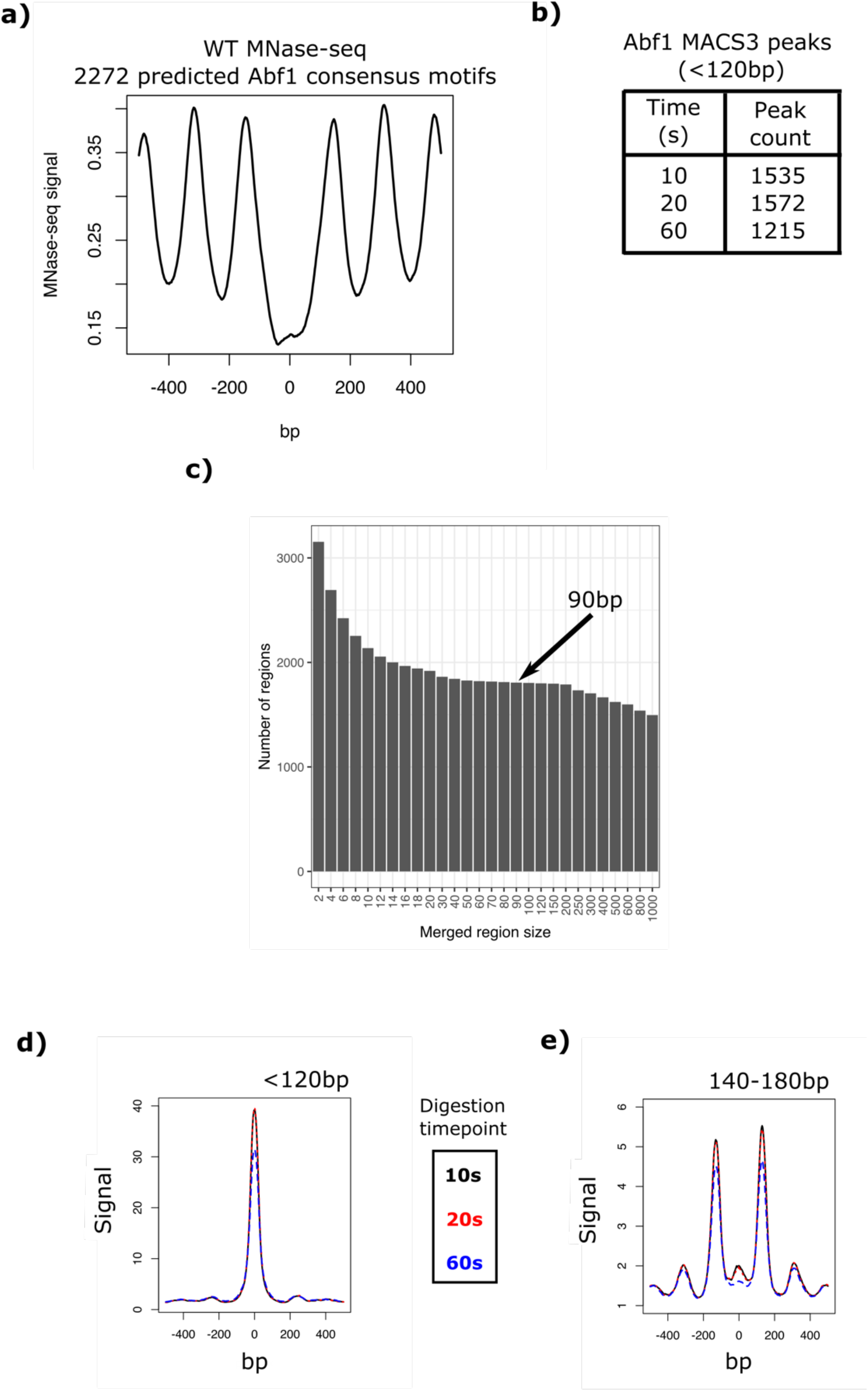
(next page) Abf1 SpLiT-ChEC signal is present at predicted sites and is found at similar sites across timepoints. a) MNase-Seq signal from wild-type yeast cells at all predicted Abf1 consensus sequences in the yeast genome. b) Counts of the number of peaks found by MACS3 at each timepoint using small-fragment signal only. c) Bar graph showing the number of peak regions reported after merging MACS3 peaks from all timepoints at increasing distances, noted on the x-axis. d) Abf1 SpLiT-ChEC SFP plotted at all center-aligned 90bp targets for each Abf1 SpLiT-ChEC timepoint. e) equivalent to d, except for Abf1 SpLiT-ChEC LFP.

**Supplemental Figure 3.**
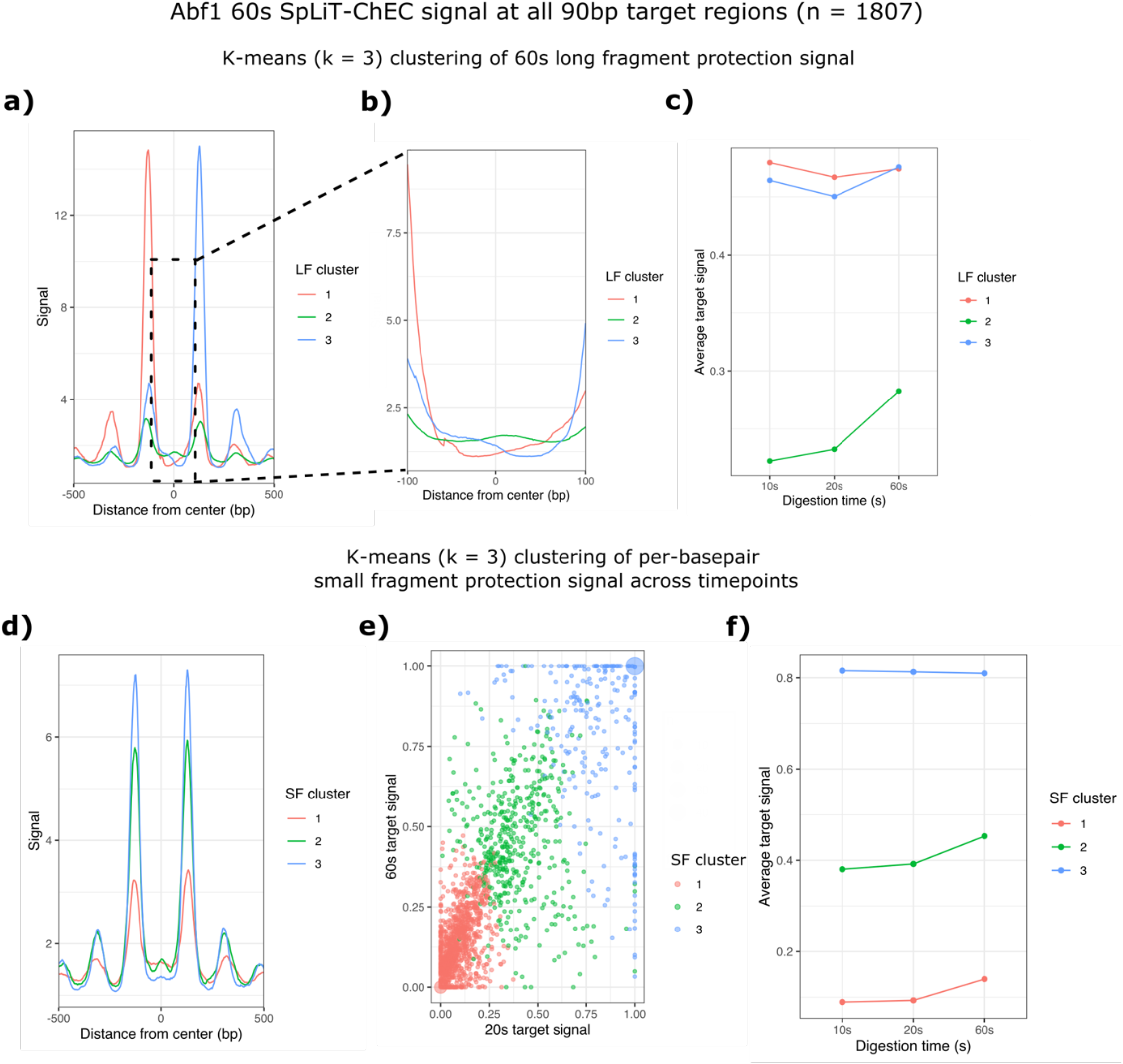
Clustering Abf1 SpLiT-ChEC signal without strand information identifies groups of similar signal. a) Average Abf1 SpLiT-ChEC LFP signal found in each cluster (k=3) after applying k-means to only the 60s LFP signal. b) Identical to a), but zoomed in on the central 200bp region of clustered signal. c) Average per-basepair signal intensity of Abf1 SpLiT-ChEC SFP signal separated by LF cluster identified in a). d) Abf1 60s SpLiT-ChEC LFP signal average traces associated with k-means clusters (k=3) obtained from clustering SFP per-basepair signal associated with each 90bp target across all three timepoints. e) Plotting 20s and 60s per-basepair signal as cartesian coordinates for each of the 90bp regions associated with SpLiT-ChEC SFP signal peaks. Coloring by SFP signal k-means cluster shows separated groups of points. f) identical to c), except the signal is grouped by SFP signal k-means clusters.

**Supplemental Figure 4.**
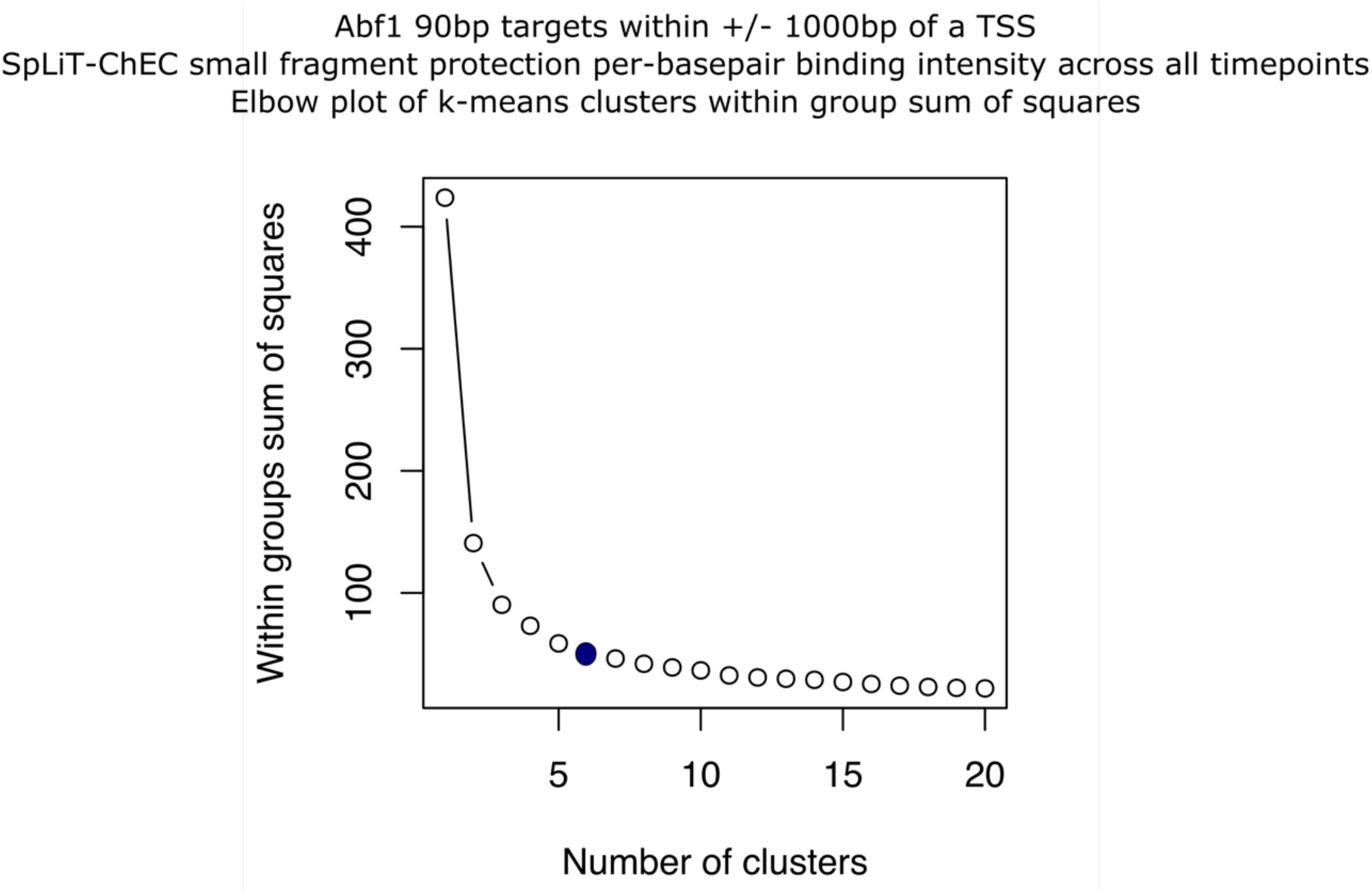
Locating optimal k-means cluster counts for three timepoints of Abf1 SFP per-basepair signal in 90bp targets near TSSs. Elbow plot of WSS calculated for each k-means cluster. The blue dot at k=6 indicates the selected cluster number.

**Supplemental Figure 5.**
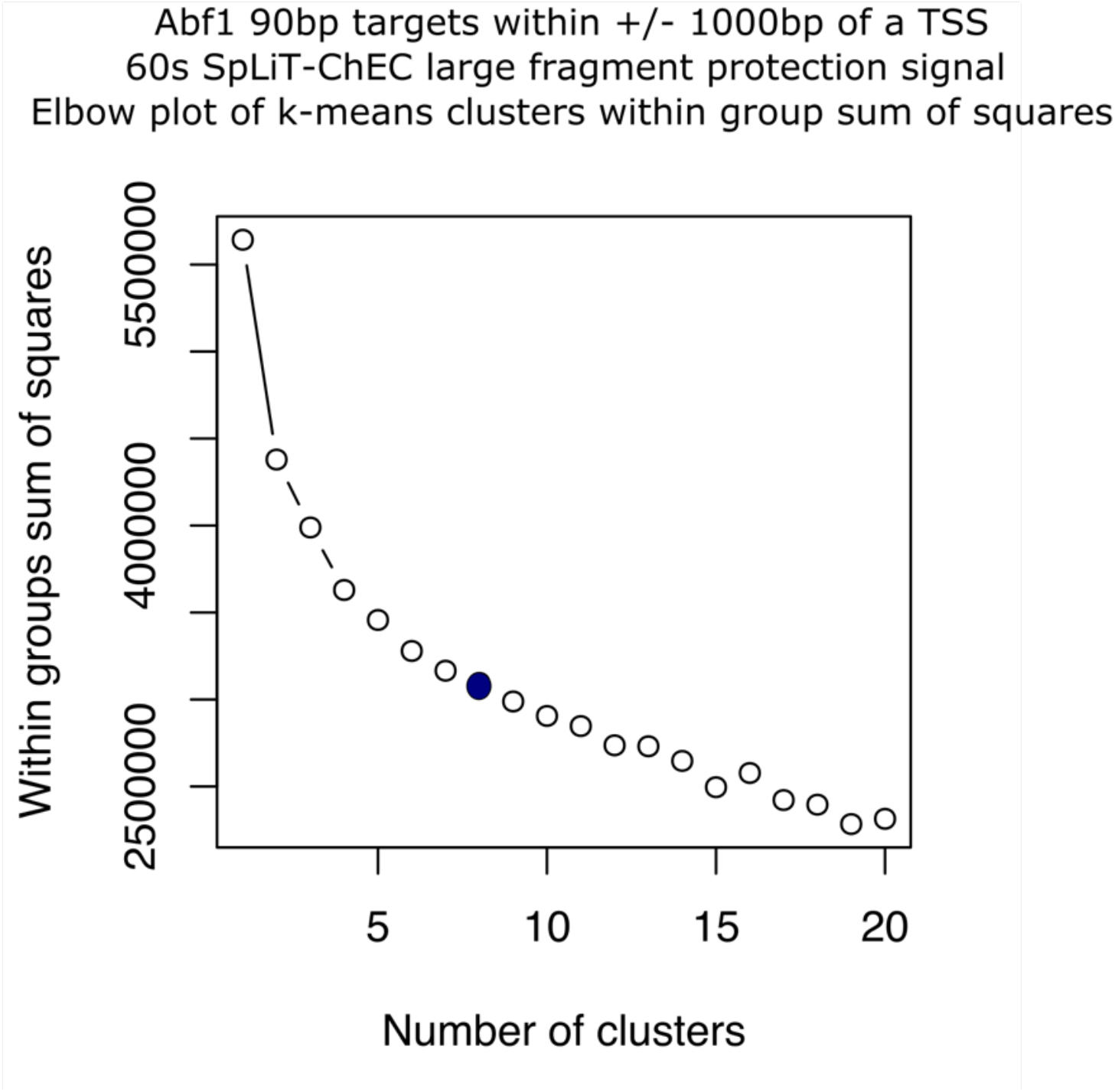
Locating optimal k-means cluster counts for Abf1 LFP signal in 90bp targets near TSSs. Elbow plot of WSS calculated for each k-means cluster. The blue dot at k=8 indicates the selected cluster number.

## References

1. Dangkulwanich M, Ishibashi T, Bintu L, Bustamante C. Molecular mechanisms of transcription through single-molecule experiments. Vol. 114, Chemical Reviews. 2014.

2. Lee TI, Rinaldi NJ, Robert F, Odom DT, Bar-Joseph Z, Gerber GK, et al. Transcriptional regulatory networks in Saccharomyces cerevisiae. Science (1979). 2002;298(5594).

3. Zamanighomi M, Lin Z, Wang Y, Jiang R, Hung Wong W. Predicting transcription factor binding motifs from DNA-binding domains, chromatin accessibility and gene expression data. Nucleic Acids Res. 2017;45(10).

4. Jolma A, Yan J, Whitington T, Toivonen J, Nitta KR, Rastas P, et al. DNA-binding specificities of human transcription factors. Cell. 2013;152(1–2).

5. Kornberg RD. Chromatin structure: A repeating unit of histones and DNA. Science (1979). 1974;184(4139).

6. Adams CC, Workman JL. Binding of disparate transcriptional activators to nucleosomal DNA is inherently cooperative. Mol Cell Biol. 1995;15(3).

7. Tsompana M, Buck MJ. Chromatin accessibility: A window into the genome. Vol. 7, Epigenetics and Chromatin. 2014.

8. Coux RX, Owens NDL, Navarro P. Chromatin accessibility and transcription factor binding through the perspective of mitosis. Transcription. 2020;11(5).

9. Clapier CR, Cairns BR. The biology of chromatin remodeling complexes. Vol. 78, Annual Review of Biochemistry. 2009.

10. Ganapathi M, Palumbo MJ, Ansari SA, He Q, Tsui K, Nislow C, et al. Extensive role of the general regulatory factors, Abf1 and Rap1, in determining genome-wide chromatin structure in budding yeast. Nucleic Acids Res. 2011;39(6).

11. Johnson DS, Mortazavi A, Myers RM, Wold B. Genome-wide mapping of in vivo protein-DNA interactions. Science (1979). 2007;316(5830).

12. Rhee HS, Pugh BF. Comprehensive genome-wide protein-DNA interactions detected at single-nucleotide resolution. Cell. 2011;147(6).

13. Kasinathan S, Orsi GA, Zentner GE, Ahmad K, Henikoff S. High-resolution mapping of transcription factor binding sites on native chromatin. Nat Methods. 2014;11(2).

14. Cui K, Zhao K. Genome-wide approaches to determining nucleosome occupancy in metazoans using MNase-Seq. Methods in Molecular Biology. 2012;833.

15. Skene PJ, Henikoff S. An efficient targeted nuclease strategy for high-resolution mapping of DNA binding sites. Elife. 2017;6.

16. Kaya-Okur HS, Wu SJ, Codomo CA, Pledger ES, Bryson TD, Henikoff JG, et al. CUT&Tag for efficient epigenomic profiling of small samples and single cells. Nat Commun. 2019;10(1).

17. Policastro RA, Zentner GE. Enzymatic methods for genome-wide profiling of protein binding sites. Brief Funct Genomics. 2018;17(2).

18. Schmid M, Durussel T, Laemmli UK. ChIC and ChEC: Genomic mapping of chromatin proteins. Mol Cell. 2004;16(1).

19. Zentner GE, Kasinathan S, Xin B, Rohs R, Henikoff S. Correction: Corrigendum: ChEC-seq kinetics discriminates transcription factor binding sites by DNA sequence and shape in vivo. Nat Commun. 2015;6(1).

20. Arnau J, Lauritzen C, Petersen GE, Pedersen J. Current strategies for the use of affinity tags and tag removal for the purification of recombinant proteins. Vol. 48, Protein Expression and Purification. 2006.

21. Zakeri B, Fierer JO, Celik E, Chittock EC, Schwarz-Linek U, Moy VT, et al. Peptide tag forming a rapid covalent bond to a protein, through engineering a bacterial adhesin. Proc Natl Acad Sci U S A. 2012;109(12).

22. Keeble AH, Banerjee A, Ferla MP, Reddington SC, Anuar INAK, Howarth M. Evolving Accelerated Amidation by SpyTag/SpyCatcher to Analyze Membrane Dynamics. Angewandte Chemie - International Edition. 2017;56(52).

23. Henikoff JG, Belsky JA, Krassovsky K, MacAlpine DM, Henikoff S. Epigenome characterization at single base-pair resolution. Proc Natl Acad Sci U S A. 2011;108(45).

24. Meers MP, Janssens DH, Henikoff S. Pioneer Factor-Nucleosome Binding Events during Differentiation Are Motif Encoded. Mol Cell. 2019;75(3).

25. Cui J, Kaandorp JA, Sloot PMA, Lloyd CM, Filatov M V. Calcium homeostasis and signaling in yeast cells and cardiac myocytes. Vol. 9, FEMS Yeast Research. 2009.

26. Chen X, Zaro JL, Shen WC. Fusion protein linkers: Property, design and functionality. Vol. 65, Advanced Drug Delivery Reviews. 2013.

27. Rhode PR, Elsasser S, Campbell JL. Role of multifunctional autonomously replicating sequence binding factor 1 in the initiation of DNA replication and transcriptional control in Saccharomyces cerevisiae. Mol Cell Biol. 1992;12(3).

28. Yarragudi A, Miyake T, Li R, Morse RH. Comparison of ABF1 and RAP1 in Chromatin Opening and Transactivator Potentiation in the Budding Yeast Saccharomyces cerevisiae. Mol Cell Biol. 2004;24(20).

29. Grant CE, Bailey TL, Noble WS. FIMO: Scanning for occurrences of a given motif. Bioinformatics. 2011;27(7).

30. Castro-Mondragon JA, Riudavets-Puig R, Rauluseviciute I, Berhanu Lemma R, Turchi L, Blanc-Mathieu R, et al. JASPAR 2022: The 9th release of the open-access database of transcription factor binding profiles. Nucleic Acids Res. 2022;50(D1).

31. McKnight LE, Crandall JG, Bailey TB, Banks OGB, Orlandi KN, Truong VN, et al. Rapid and inexpensive preparation of genome-wide nucleosome footprints from model and non-model organisms. STAR Protoc. 2021;2(2).

32. Bailey TL, Grant CE. SEA: Simple Enrichment Analysis of motifs. bioRxiv. 2021

33. Grant CE, Bailey TL. XSTREME: Comprehensive motif analysis of biological sequence datasets. bioRxiv. 2021

34. O Connor T, Grant CE, Boden M, Bailey TL. T-Gene: Improved target gene prediction. Bioinformatics. 2020;36(12).

35. Yen K, Vinayachandran V, Batta K, Koerber RT, Pugh BF. Genome-wide nucleosome specificity and directionality of chromatin remodelers. Cell. 2012 Jun 22;149(7):1461–73.

36. Ganguli D, Chereji R V., Iben JR, Cole HA, Clark DJ. RSC-dependent constructive and destructive interference between opposing arrays of phased nucleosomes in yeast. Genome Res. 2014;24(10).

37. Weiner A, Hughes A, Yassour M, Rando OJ, Friedman N. High-resolution nucleosome mapping reveals transcription-dependent promoter packaging. Genome Res. 2010;20(1).

38. Balsalobre A, Drouin J. Pioneer factors as master regulators of the epigenome and cell fate. Vol. 23, Nature Reviews Molecular Cell Biology. 2022.

39. Kubik S, O’Duibhir E, de Jonge WJ, Mattarocci S, Albert B, Falcone JL, et al. Sequence-Directed Action of RSC Remodeler and General Regulatory Factors Modulates +1 Nucleosome Position to Facilitate Transcription. Mol Cell. 2018;71(1).

40. Barnes T, Korber P. The active mechanism of nucleosome depletion by Poly(da:dt) tracts in vivo. Int J Mol Sci. 2021;22(15).

41. Ju QD, Morrow BE, Warner JR. REB1, a yeast DNA-binding protein with many targets, is essential for growth and bears some resemblance to the oncogene myb. Mol Cell Biol. 1990;10(10).

42. Gibson DG, Young L, Chuang RY, Venter JC, Hutchison CA, Smith HO. Enzymatic assembly of DNA molecules up to several hundred kilobases. Nat Methods. 2009;6(5).

43. Walker A, Taylor J, Rowe D, Summers D. A method for generating sticky-end PCR products which facilitates unidirectional cloning and the one-step assembly of complex DNA constructs. Plasmid. 2008;59(3).

44. Andrews S, others. FastQC: a quality control tool for high throughput sequence data. 2010. Https://Www.BioinformaticsBabrahamAcUk/Projects/Fastqc/. 2019;

45. Howe KL, Achuthan P, Allen J, Allen J, Alvarez-Jarreta J, Ridwan Amode M, et al. Ensembl 2021. Nucleic Acids Res. 2021;49(D1).

46. Langmead B, Salzberg SL. Fast gapped-read alignment with Bowtie 2. Nat Methods. 2012;9(4).

47. Li H, Handsaker B, Wysoker A, Fennell T, Ruan J, Homer N, et al. The Sequence Alignment/Map format and SAMtools. Bioinformatics. 2009;25(16).

48. Ramírez F, Ryan DP, Grüning B, Bhardwaj V, Kilpert F, Richter AS, et al. deepTools2: a next generation web server for deep-sequencing data analysis. Nucleic Acids Res. 2016;44(W1).

49. Quinlan AR, Hall IM. BEDTools: A flexible suite of utilities for comparing genomic features. Bioinformatics. 2010;26(6).

